# PLCβs are recruited to the plasma membrane in macrophages by both Gβγ and Gα_q_

**DOI:** 10.64898/2026.01.28.702352

**Authors:** Maria E. Falzone, Priyam Banerjee, Roderick MacKinnon

**Affiliations:** Laboratory of Molecular Neurobiology and Biophysics, The Rockefeller University, NY, United States; Howard Hughes Medical Institute, The Rockefeller University, NY, United States; Department of Biochemistry and Structural Biology and Greehey Children’s Cancer Research Institute, University of Texas Health Science Center at San Antonio, TX, United States; Bio-Imaging Resource Center, The Rockefeller University, NY, United States

**Keywords:** PLCβ3, PIP2, Gβγ, GPCR signaling, Macrophages

## Abstract

PLCβ enzymes cleave PIP2 from the plasma membrane, producing IP3 and DAG, which regulate intracellular Ca^2+^ levels and protein kinase C activity, respectively. They are regulated by GPCR signaling through the G proteins Gβγ and Gα_q_ and have been shown to function as coincidence detectors for dual stimulation of Gα_q_ and Gα_i_-coupled receptors via these G proteins. PLCβs are aqueous-soluble enzymes, but partition onto the membrane surface to access their lipid substrate. We previously demonstrated that membrane recruitment and orientation of the catalytic core on the membrane surface underlie Gβγ-dependent regulation of PLCβ enzymes. Using macrophages as a model system, where PLCβ signaling is essential for responses to infection and tissue injury, we investigated the contribution of Gβγ-dependent regulation and membrane recruitment of PLCβ in the context of endogenous signaling. By measuring Ca^2+^ mobilization, we demonstrate that both Gα_i_ and Gα_q_-coupled receptors independently stimulate PLCβ activity. Using total internal reflection and stimulated emission depletion microscopy, we demonstrate that most of the PLCβ3 in the cell is localized away from the plasma membrane at rest but is rapidly recruited to the plasma membrane upon stimulation by both Gα_i_ and Gα_q_-coupled receptors, illustrating that both Gβγ and Gα_q_ recruit PLCβ to the plasma membrane. These results support an updated model for G protein-dependent regulation of PLCβ enzymes, where Gβγ-induced regulation in the absence of Gα_q_ can occur and is apparently dictated by the local concentration of receptor, G proteins, and PLCβ.

**Significance Statement:** PLCβ enzymes are critical mediators of signal transduction with roles in neuronal, cardiac, and immunological signaling. Despite this importance, many aspects of their function and regulation remain poorly understood. PLCβs are aqueous soluble but must partition onto the membrane surface to access their lipid substrate, which enables regulation at the partitioning step, the catalytic step, or both. We previously demonstrated that membrane recruitment and orientation of the catalytic core on the membrane surface underlie PLCβ regulation by Gβγ in a reconstituted system. Using macrophages as a model system for physiological signaling, we demonstrate here that Gβγ can activate PLCβ via membrane recruitment in cells.

## Introduction

PLCβ enzymes are canonical downstream effectors of G protein coupled receptor (GPCR) signaling (1-3). Upon activation, PLCβ enzymes cleave PIP2 from the plasma membrane, producing inositol triphosphate (IP3) and DAG (4, 5). IP3 opens the IP3 receptor ion channel on the endoplasmic reticulum (ER), leading to an increase in cytosolic Ca^2+^ concentration and DAG activates protein kinase C (PKC). PLCβ function is essential for a multitude of cellular processes, including cardiac signaling, neuronal signaling, and macrophage activation (1-3). Accordingly, their activity needs to be tightly regulated, with very low basal activity and robust stimulus-dependent activation. PLCβs are under the control of both Gα_q_ and Gα_i_-dependent GPCR signaling through direct interactions with the G proteins Gα_q_ and Gβγ (1, 3). They have also been shown to function as coincidence detectors for co-stimulation of Gα_q_ and Gα_i_-coupled receptors through simultaneous activation by both G proteins (1-3).

Using *in vitro* reconstitution experiments, we previously demonstrated that Gβγ and Gα_q_ activate PLCβ via differing mechanisms, Gβγ by recruiting PLCβ to the membrane and orienting the catalytic core for substrate binding and catalysis and Gα_q_ by increasing the catalytic rate constant, k_cat_, by displacing an autoinhibitory loop from the active site (6, 7). Additionally, we showed that dual stimulation by both G proteins is mediated by the combined effects of these differing activation mechanisms. Thus, together, both G proteins activate to the order of the product of each individual effect (6). These observations led us to propose that a combination of autoinhibition and robust stimulus-dependent activation allow PLCβ enzymes to be tightly regulated by GPCR signaling and prevent aberrant signaling.

While we and others have demonstrated that Gβγ robustly activates PLCβ enzymes in reconstituted systems (7-11), the role and feasibility of independent regulation of PLCβ enzymes by Gβγ in the cellular context has come under debate. Pertussis toxin (PTX)-sensitive and Gβγ-dependent regulation of PLCβs has been reported for decades (1, 3), leading to the proposal that Gβγ released from Gα_i_-coupled receptor signaling could activate PLCβs independently of Gα_q_ (12, 13). However, several recent reports have proposed that the presence of Gα_q_ is strictly required for any Gβγ-dependent activation (14-19). Macrophages are one cellular system that have been shown to support both Gα_i_ and Gα_q_-dependent PLCβ activity, where both pathways play important roles in macrophage activation in response to cellular injury and infection (12, 20-22). To understand how G protein regulation of PLCβs is manifested in the native cellular context, and whether each pathway can separately influence PLCβ activity, we studied this cellular system.

Using Raw264.7 cells, a mouse macrophage cell line with endogenous Gα_i_ and Gα_q_-dependent PLCβ activation and coincidence detection, we investigated the regulation of PLCβ activity and the role of membrane recruitment in this regulation. Importantly, we used the endogenous signaling components without overexpression to study signaling in the native context. Our results show that Gα_i_-coupled signaling is sufficient to activate PLCβ robustly in the absence of Gα_q_ stimulation and appears to do so by recruiting PLCβ to the plasma membrane on a functionally relevant timescale. These observations lead us to propose that Gα_i_ signaling can activate PLCβ enzymes in Raw264.7 cells by membrane recruitment. The ability of Gα_i_ signaling to activate PLCβ in these cells likely depends on the concentration and distribution of receptor, G proteins, and PLCβ.

## Results

### Stimulation of Raw264.7 cells with GPCR agonists results in PLCβ-dependent Ca^2+^ transients

To confirm the presence of GPCR-dependent PLCβ activity in Raw264.7 cells, we measured PLCβ-dependent increases in cytoplasmic Ca^2+^ concentration using a fluorescent Ca^2+^ indicator. Wildtype Raw264.7 cells were loaded with Fluo-4 AM (**Fig. 1A**), followed by subsequent stimulation with agonists for three different receptors: a complement receptor (C5aR), a purinergic receptor (P2Y6R), and a lysophosphatidic acid receptor (LPAR). The agonist C5a activates C5aR, which is part of the complement pathway and plays a role in macrophage activation and response to infection and predominantly signals through Gα_i_ (23). The agonist UDP signals through P2Y6R and plays a role in macrophage activation in response to tissue injury and predominantly signals through Gα_q_ (24). The agonist LPA signals via LPAR and plays a role in macrophage activation and inflammation and predominantly signals through Gα_q_ (25). In agreement with previous results, addition of all three agonists lead to rapid Ca^2+^ transients, which peak ∼10s following stimulation and decay to baseline over 40-100s (**Fig. 1B, S1A**) (12, 20, 26). Treatment with PLCβ inhibitor U73122 blocked these Ca^2+^-transients, whereas vehicle (DMSO) and U73342, the established negative control compound without PLCβ inhibitory activity, had only small reductions in peak height and slope, indicating these transients are PLCβ-dependent (**Fig. 1C, S1**).

**Figure 1:**
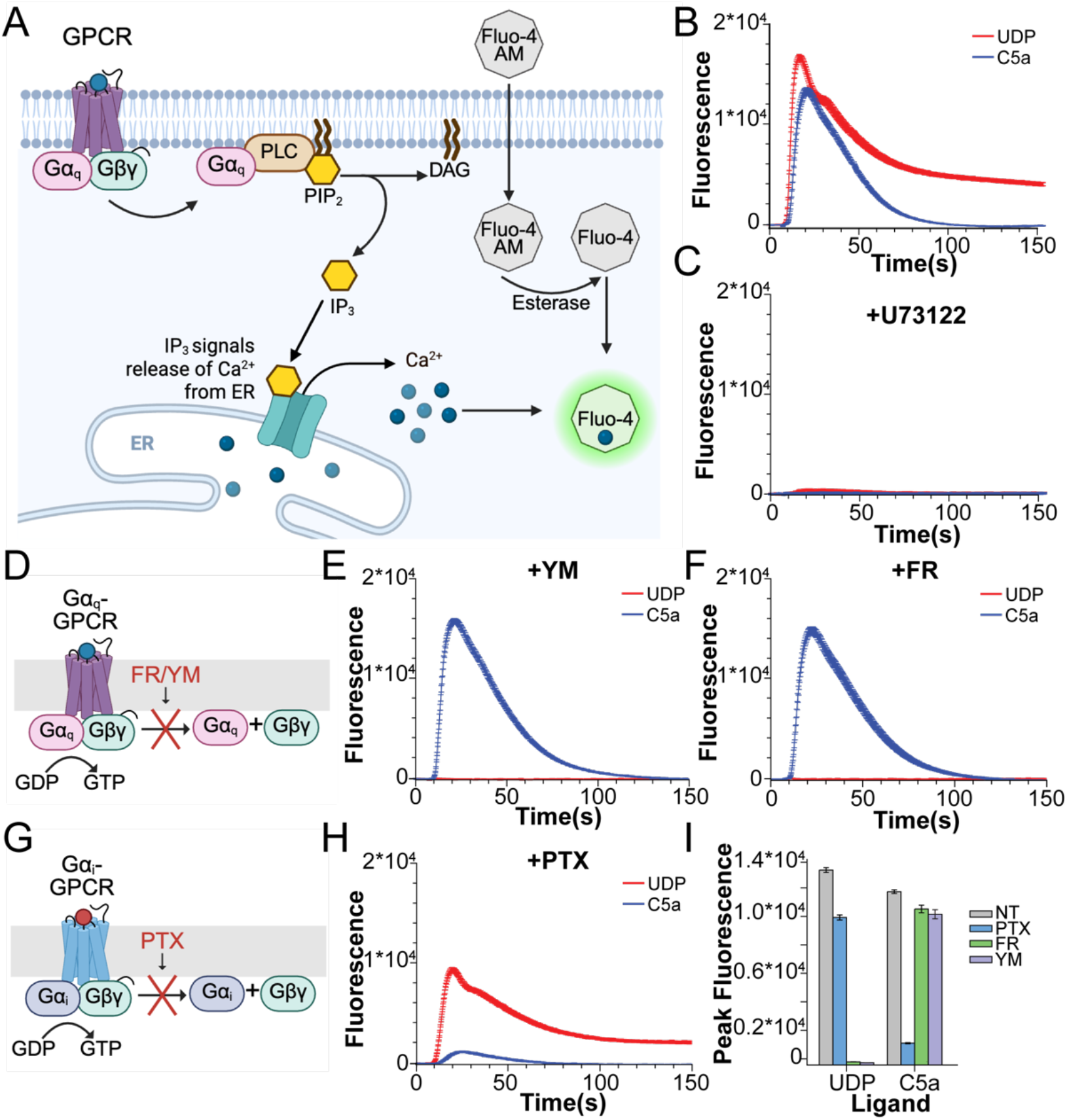
PLCβ-dependent signaling in Raw264.7 cells. **A**: Cartoon of fluo-4 Ca^2+^ assay for PLCβ activity. Fluo-4 AM is loaded into cells and esterases cleave the hydrophobic groups, trapping it inside. GPCR stimulation activates PLCβ, which generates IP3, resulting in Ca^2+^ release from the ER through the IP3 receptor and increases the fluorescence signal. **B-C**: Representative Ca^2+^ transients upon stimulation with UDP (red) and C5a (blue) in the absence (B) or presence (C) of PLCβ inhibitor U73122 (20 μM). **D**: Cartoon representation of the effects of Gα_q_ cyclic peptide inhibitors FR/YM. **E-F**: Representative Ca^2+^ transients upon stimulation with UDP (red) and C5a (blue) in the presence of Gα_q_ inhibitors FR (10 μM) (E) or YM (500 nM) (F). **G**: Cartoon representation of the effects of Gα_i_ signaling inhibitor PTX. **H**: Representative Ca^2+^ transients upon stimulation with UDP (red) and C5a (blue) in the presence of 300 ng/mL PTX. **I**: Quantification of Ca^2+^ peak fluorescence for UDP and C5a in the presence of PTX, FR, or YM. Error bars are SEM. For B-C, E-F, and H, traces are averaged from 10 wells and error bars are SEM. UDP was added at 10 μM and C5a at 20 nM final concentrations.

### G protein coupling of P2Y6R, C5aR, and LPAR

Many GPCRs couple to multiple G proteins. In order to assess the role of Gα_i_ versus Gα_q_-coupling in regulation of PLCβ enzymes, we sought to define the G protein coupling of our three receptors using FR900359 (FR) and YM254890 (YM), cyclic peptide inhibitors that selectively block Gα_q_βγ trimer separation (27), and pertussis toxin (PTX), which selectively blocks Gα_i_βγ trimer separation (28) (**Fig. 1D, G**). Blocking Gα_q_ signaling completely inhibited UDP and LPA-dependent Ca^2+^ increases, whereas blocking Gα_i_ signaling inhibited most of the C5a-dependent Ca^2+^ increase with minimal effect on the UDP-dependent increase (**Fig. 1D-I, S2A-C**). These observations indicate that UDP and LPA signal through Gα_q_ and C5a signals through Gα_i_ in this cell line, consistent with previous results (12, 20, 21). Small decreases in UDP and LPA-induced Ca^2+^ peak height and slope in the presence of PTX (**Fig. 1H-I, S2A-C**) are likely due to non-specific toxicity, since their effects are completely blocked by FR/YM treatment, indicating they are completely dependent on Gα_q_. Importantly, C5a-dependent Ca^2+^ transients are intact upon FM and YR treatment, demonstrating that Gα_q_ is not required for Gα_i_-dependent PLCβ activation in macrophages, consistent with previous results (12).

### Dual stimulation of C5aR and P2Y6

We next sought to evaluate the effect of dual stimulation of Gα_i_- and Gα_q_-coupled receptors on PLCβ activation. We stimulated cells with a combination of C5a and UDP and measured the corresponding Ca^2+^ increases. While the peak fluorescence did not increase on the order of the sum of each agonist alone, the transients arose significantly faster than in the presence of each ligand on its own (**Fig. 2A-B**). To quantify the rate of transient increase, we fit the early rise of the Ca^2+^ signal to a straight line and compared the slopes (**Fig. 2B-C**). The slope upon dual stimulation is significantly greater than the sum of the slopes of the individual stimuli (**Fig. 2C**), suggesting that the largest effect of dual stimulation is a faster Ca^2+^ increase. Using the slope as a measure of PLCβ activation, dual stimulation of Gα_i_ and Gα_q_-coupled receptors yields approximately the product of each ligand on its own, which is consistent with previous work and our *in vitro* results (6, 12). Additionally, upon inhibition of Gα_i_ or Gα_q_-dependent signaling using PTX or FM/YR, the slopes of the Ca^2+^ increases return to the level of each agonist on its own, pointing to the absence of additional synergy in this regulation (**Fig. S2D-F**).

**Figure 2:**
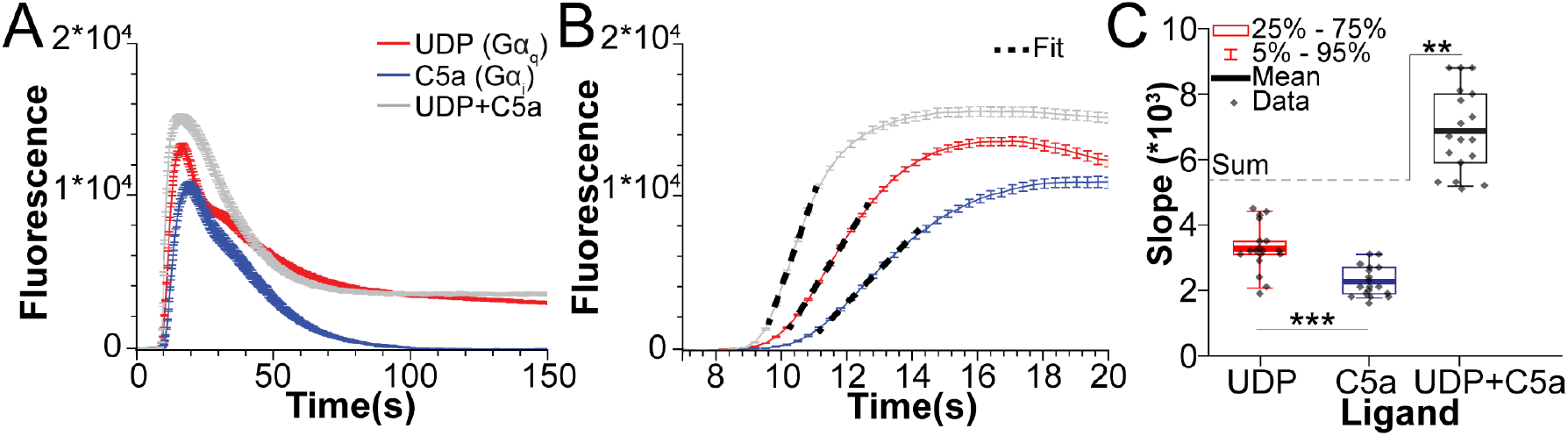
Effects of dual stimulation of Gα_i_ and Gα_q_-coupled receptors on PLCβ activity. **A-B**: Representative Ca^2+^ transients upon stimulation with UDP (red), C5a (blue), or UDP+C5a (gray) for the complete time course (A) or for early times (B). Traces are averaged from 10 wells and error bars are SEM. UDP was added at 10 μM and C5a at 20 nM final concentrations. Fits of the increase to a straight line are shown as a black dashed line in B. **C**: Slopes from the fits in B. The average of the sums of UDP and C5a alone is denoted with a gray dashed line. *: 0.05>p>0.005. **: 0.005>p>0.0005, ***: p<0.0005.

### Both Gα_q_ and Gα_i_-dependent signaling recruit PLCβ3 to the plasma membrane

We next studied the PLCβ cellular distribution and how it is influenced by GPCR stimulation using immunolabeling with primary and secondary antibodies. We used a primary antibody specific for mouse PLCβ3, which has been shown to underlie coincidence detection in Raw264.7 cells (12), and mouse C5aR to serve as a plasma membrane marker. We investigated the cellular distribution of PLCβ3 and its recruitment to the plasma membrane using three different types of microscopy to ensure that our observations were robust and to increase the confidence in our conclusions. (**Fig. S3A**).

To get a broad overview of the PLCβ3 present on the plasma membrane and how it is influenced by agonist stimulation, we used Total Internal Reflection (TIRF) microscopy, with a penetration depth of ∼0.100 μm (29) (**Fig. S3A, C**). In the absence of stimulation, we observed PLCβ3 puncta with a mean density of ∼0.2/μm^2^ and a spot area of ∼0.13 μm^2^. This spot area is ∼2-fold larger than expected for the diffraction limited spot diameter of 0.250 μm (∼0.05 μm^2^), which could be due to clusters of several PLCβ3 proteins. Upon stimulation for 3-120s with UDP, C5a, or both, we observed a significant increase in PLCβ3 spot density and size (**Fig. 3**), suggesting PLCβ3 is recruited to the plasma membrane upon receptor activation. Stimulation with C5a significantly increased PLCβ3 spot density after 5s and spot size after 10s, with a maximal increase of ∼3-fold for both size and density after 120s (**Fig. 3F, I**). Stimulation with UDP significantly increased PLCβ3 spot density after 10s and spot size after 5s, with a maximal increase of 2-fold for both spot size and density after 60s (**Fig. 3G, J**). Stimulation with both agonists significantly increased PLCβ3 spot size and density at 5s, with a maximal effect of ∼4-fold for density and ∼3-fold for size after 120s (**Fig. 3H, K**), consistent with our functional experiments in that the greatest effect of dual stimulation is a faster response. Otherwise, there is no significant difference between dual stimulation and C5a stimulation alone. To ensure that our observations of membrane recruitment correlated with functional measurements of Ca^2+^ signals, we selectively disrupted Gα_i_ and Gα_q_-coupled signaling and assessed the effect on C5a-induced membrane recruitment (Fig S4). Upon disruption of Gα_i_-coupled signaling with PTX, PLCβ membrane recruitment is abolished up to 30s following C5a stimulation. In contrast, we observe a decrease in PLCβ puncta number and size at the plasma membrane with C5a stimulation (Fig S4 A-D), which we attribute to toxicity associated with PTX treatment because the cells appeared visibly less healthy. Upon disruption of Gα_q_-coupled signaling with FR, C5a-induced PLCβ membrane recruitment is comparable to cells in the absence of FR, with significant increases in PLCβ puncta number and size at 10s and 30s (Fig. S4 E-G). Together, these studies support the proposal that free Gβγ from Gα_i_-coupled receptors is sufficient to recruit PLCβ to the membrane and drive signaling.

**Figure 3:**
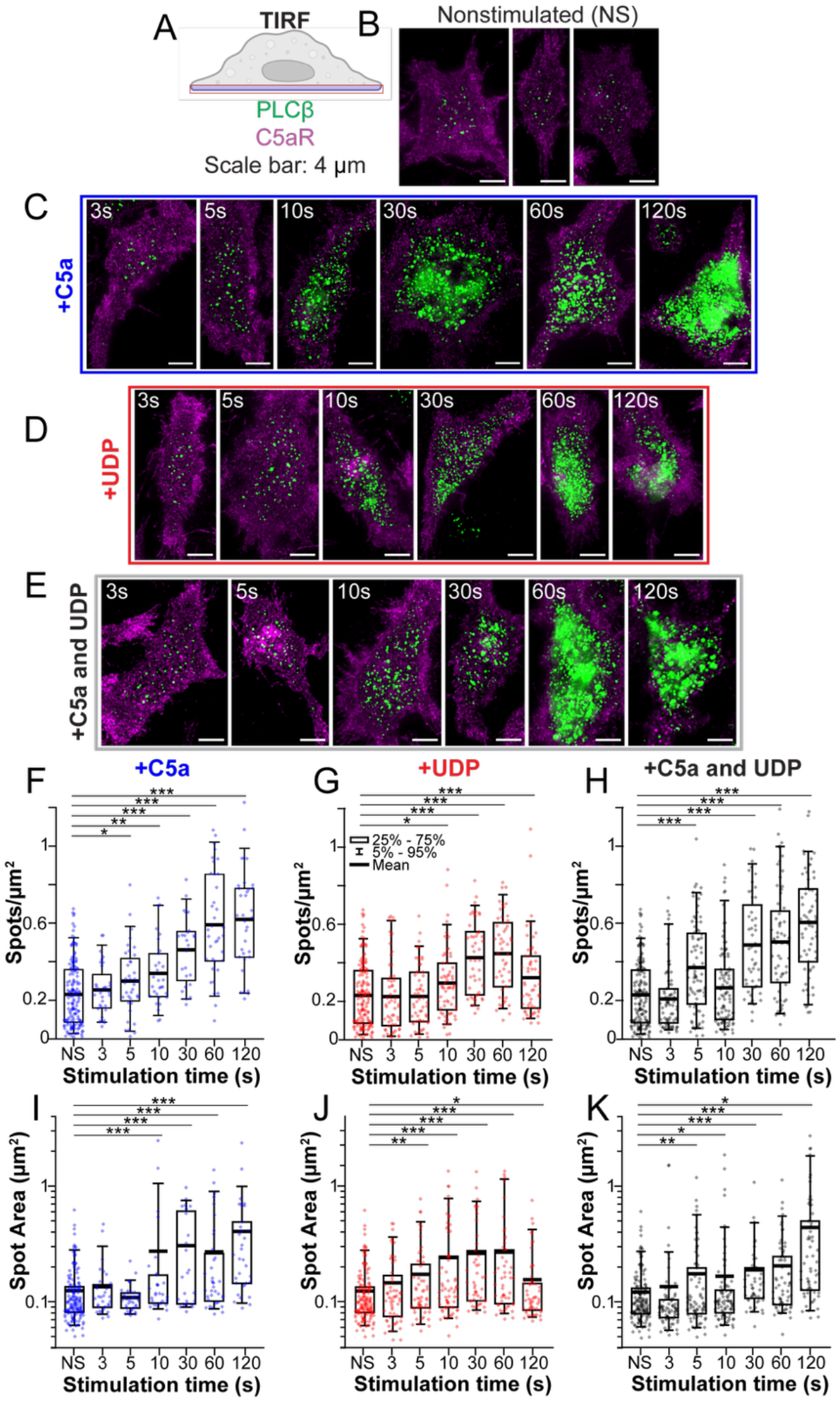
GPCR signaling recruits PLCβ3 to the plasma membrane. **A**: Cartoon of the TIRF imaging plane. **B-E**: Representative images of cells stained for PLCβ3 (green) or C5aR (purple) as a membrane marker in the absence of stimulation (B) or following stimulation with 20 nM C5a (C), 10 μM UDP (D), or UDP+C5a (E). Scale bar is 4 μm. Time points indicate the time of agonist exposure prior to cell fixation. **F-H**: Quantification of PLCβ3 spot density over time upon stimulation with C5a (F), UDP (G), or UDP+C5a (H). **I-K**: Quantification of PLCβ3 spot size over time upon stimulation with C5a (I), UDP (J), or UDP+C5a (K). NS is nonstimulated. Each point is a single cell. *: 0.05>p>0.005. **: 0.005>p>0.0005, ***: p<0.0005.

In our functional experiments, the Ca^2+^ transient peaks ∼10s after stimulation and then decays over 40-100s. In contrast, PLCβ3 puncta at the plasma membrane continue increasing in size and density as stimulation time increases up to 120s. This discrepancy is likely due to tight cellular control of cytoplasmic Ca^2+^ concentration, the readout used in our functional assay, as it is likely pumped back into the ER despite continued stimulation. Alternatively, the short timescale of our functional signal could be due to depletion of PIP2 from the plasma membrane. Accordingly, we focus on PLCβ3 function within 10s of stimulation (**Fig. S3B**), what we consider to be a functionally relevant timescale for signaling. Our data suggest that both UDP and C5a recruit PLCβ3 to the plasma membrane on this timescale.

### PLCβ is mostly localized to the interior of the cell at rest

To gain information about PLCβ3 distribution across the cell volume, we carried out 3D Stimulated Emission Depletion (STED) microscopy axial line scans on Raw264.7 cells fixed and stained for PLCβ3 and C5aR (**Fig. 4, S3A**) (30). A major advantage of the 3D STED technique is its ability to resolve structures at isotropic (X,Y,Z) super-resolution beyond the diffraction limit, down to ∼80-100 nm (**Fig. S3A**) (30). We used the C5aR signal to trace the plasma membrane and quantified the percent of PLCβ3 puncta within 100 nm, corresponding to the resolution limit. In the absence of stimulation, PLCβ3 is predominately localized in the interior of the cell, with only ∼20% of puncta at the plasma membrane (**Fig. 4**). Upon stimulation with UDP, C5a, or both, the percentage of PLCβ3 puncta at the plasma membrane increases, consistent with the results observed with TIRF microscopy. However, only C5a and the combination of both agonists have a significant effect over the functionally relevant timescale of 10s (**Fig. 4**). Stimulation with UDP increases the percentage of PLCβ3 puncta at the plasma membrane ∼1.3-fold after 60s of stimulation (**Fig. 4G**). Stimulation with C5a and both agonists increase the percentage of puncta of PLCβ3 at the plasma membrane ∼1.7-fold after 60s of stimulation (**Fig. 4F, H**). While the outcome of 3D STED shows a statistically significant trend over a limited range, we investigate membrane partitioning of PLCβ next using a different approach.

**Figure 4:**
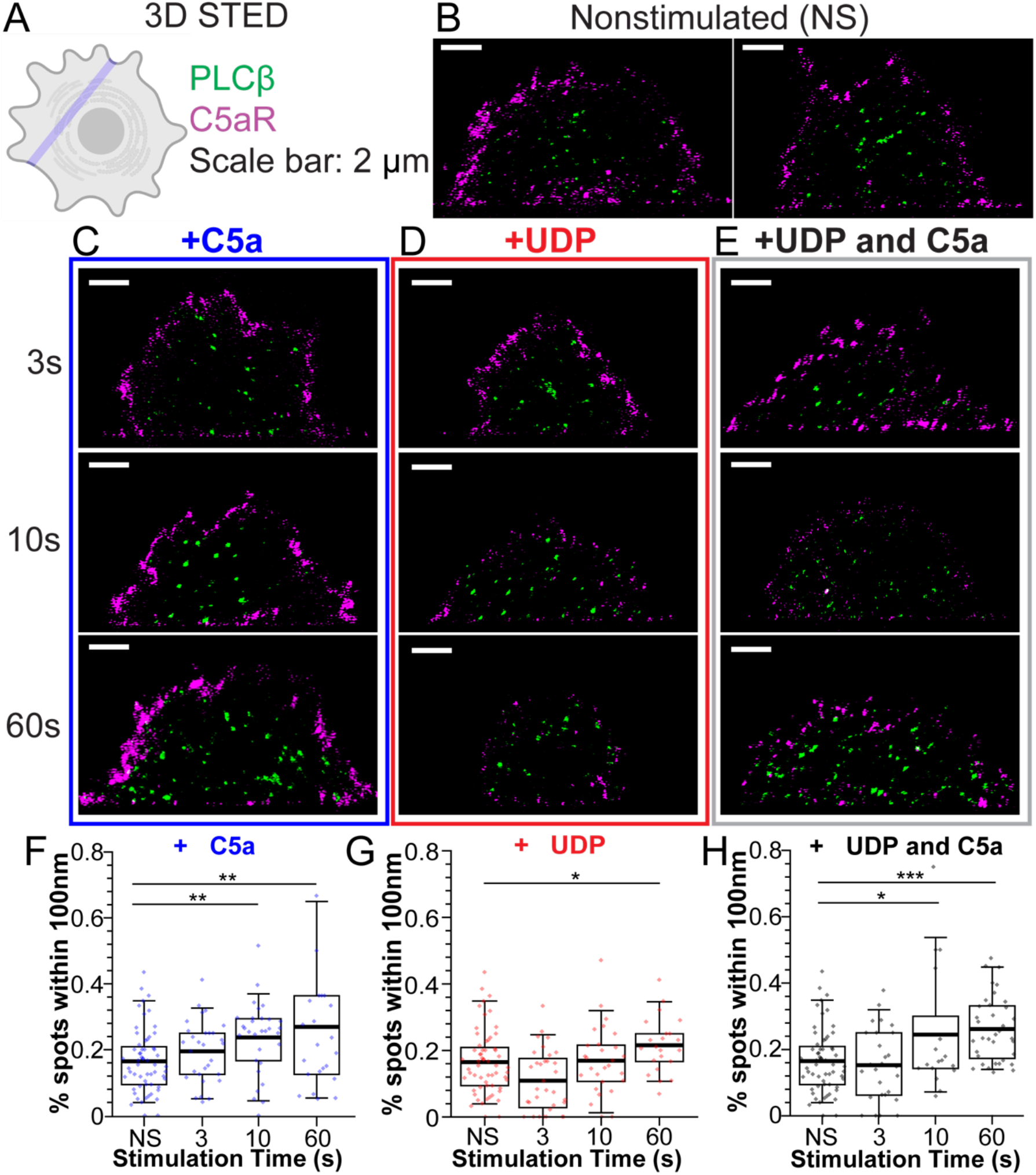
The majority of PLCβ3 is localized in the interior of the cell at rest. **A**: Cartoon of 3D STED linescan imaging plane. **B-E**: Representative images of cells stained for PLCβ3 (green) or C5aR (purple) as a membrane marker in the absence of stimulation (B) or following stimulation with 20 nM C5a (C), 10 μM UDP (D), or UDP+C5a (E). Scale bar is 2 μm. Time points indicate the time of agonist exposure prior to cell fixation. **F-H**: Quantification of percentage of PLCβ3 spots within 100 nm of the plasma membrane over time upon stimulation with C5a (F), UDP (G), or UDP+C5a (H). NS is nonstimulated. Boxes are 25%-75%, error bars are 5%-95%, thick bar is the mean. Each point is a single linescan. *: 0.05>p>0.005. **: 0.005>p>0.0005, ***: p<0.0005.

### Gα_i_-coupled signaling is more efficient at recruiting PLCβ3 to the plasma membrane at short time scales

To obtain information about the PLCβ3 distribution on the plasma membrane in a different way, we used super-resolution 2D STED imaging, which permits higher 2D (X,Y) resolution (∼40-50 nm) (**Fig. S3**) (30), allowing for investigation of the diffraction-limited puncta observed in the TIRF experiments. We collected z-stacks of the plasma membrane of Raw264.7 cells fixed and stained for PLCβ3 and C5aR, guided by the C5aR signal as a plasma membrane marker. In the absence of stimulation, the PLCβ3 density was ∼4 puncta/μm^3^ with an average puncta lateral diameter of ∼70 nm, suggesting that the spots observed in the TIRF imaging are sometimes due to multiple PLCβ3s within the 250 nm diffraction-limited resolution. We observe clear recruitment of PLCβ3 to the plasma membrane upon stimulation with Gα_i_ or Gα_q_-coupled agonists. Upon stimulation with C5a, the PLCβ3 puncta density is increased by ∼2.5-fold after 10s, (**Fig. 5C-E**). Upon stimulation with UDP, the PLCβ3 puncta density is increased by ∼2.5-fold after 60s but not after 10s, consistent with the 3D STED measurements (**Fig. 5F-H**) Finally, upon stimulation with both agonists, the PLCβ3 puncta density is increased ∼2.5-fold at 10s and ∼3-fold at 60s (**Fig. 5I-K**). Across all stimulation conditions, the PLCβ3 X-Y diameter did not change significantly.

**Figure 5:**
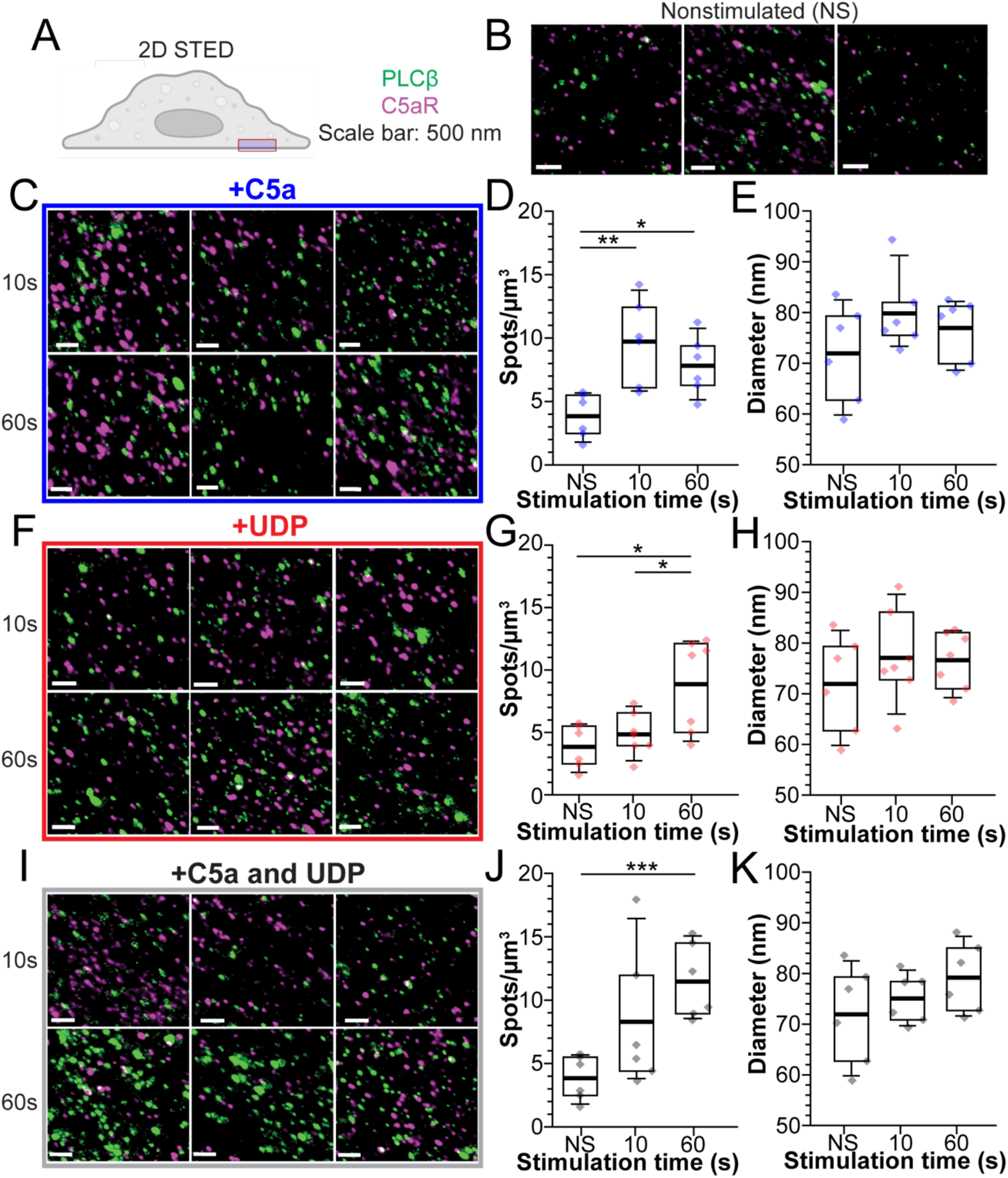
2D STED demonstration of GPCR-dependent PLCβ3 membrane recruitment. **A**: Cartoon of the 2D STED imaging plane. **B,C,F,I**: Representative images of cells stained for PLCβ3 (green) or C5aR (purple) as a membrane marker in the absence of stimulation (B) or following stimulation with 20 nM C5a (C), 10 μM UDP (D), or UDP+C5a (E). Scale bar is 500 nm. Time points indicate the time of agonist exposure prior to cell fixation. Images are maximum intensity projections of Z stacks for representative purposes. **D,G,J**: PLCβ spot density on the plasma membrane over time upon stimulation with C5a (D), UDP (G), or UDP+C5a (J). **E,H,K**: Quantification of PLCβ3 X-Y diameter over time upon stimulation with C5a (E), UDP (H), or UDP+C5a (K). NS is nonstimulated. Boxes are 25%-75%, error bars are 5%-95%, thick bar is the mean. Each point is a single z stack. *: 0.05>p>0.005. **: 0.005>p>0.0005, ***: p<0.0005.

Quantification of object number and geometry in light microscopy approaches is dependent on the signal threshold used for segmentation. To reduce bias, we completed the analysis of our 2D STED data at a low segmentation threshold **(Fig. S5)**. With low thresholds, the density of PLCβ3 on the plasma membrane was ∼1.7 puncta/μm^3^ with an X-Y diameter of ∼80 nm. The increases in PLCβ3 density at the plasma membrane upon stimulation with C5a, UDP, or both were consistent with the higher segmentation thresholds, suggesting our measurements scaled proportionately with signal thresholds, and that our conclusions are robust to variations in signal threshold over the tested range. Given the improved resolution, these observations suggest that Gα_i_-coupled signaling may be more efficient at recruiting PLCβ to the plasma membrane than Gα_q_-coupled signaling on a functionally relevant time scale.

### C5aR density at the plasma membrane increases upon C5a stimulation

C5aR was used as a plasma membrane marker in our 2D STED experiments, which enabled analysis of its puncta density and size. At rest, the C5aR density was ∼10 puncta/μm^3^ with an X-Y diameter of ∼120 nm, consisting of larger and more concentrated puncta than PLCβ (**Fig. S6**). Upon stimulation with C5a, we observed the density and diameter of C5aR puncta increased ∼1.5-fold (**Fig. S6**). This effect was not observed upon stimulation with an unrelated agonist, UDP. In the presence of both agonists, the increases are evident but smaller. These observations suggest that receptor mobilization may play a role in cellular responses to C5a stimulation.

## Discussion

This study aims to elucidate the role and mechanism of Gβγ-dependent activation of PLCβ enzymes in the cellular context. We previously demonstrated that Gβγ robustly activates PLCβ via membrane recruitment and orientation using *in vitro* reconstitution studies (6, 7). However, the question of whether Gβγ recruits PLCβ to the membrane and its ability to independently regulate PLCβ in the cellular environment is debated (14-19). To address this question, we used a mouse macrophage cell line, Raw264.7 cells, to study GPCR-dependent regulation of PLCβ function and cellular localization. Using endogenous proteins to observe native signaling, our experiments show that stimulation of both Gα_q_ and Gα_i_-coupled receptors induce PLCβ-dependent Ca^2+^ transients (**Fig. 1-2**). Gα_i_-dependent signaling was intact upon treatment with FR/YM, cyclic peptide inhibitors that block Gα_q_βγ trimer dissociation, demonstrating that Gα_q_ is dispensable for Gα_i_-dependent PLCβ activation in this context, in agreement with previous results (12) (**Fig. 1**). One potential caveat to these observations is that the Gα_q_-like G protein Gα_15_ is expressed in macrophages and has been shown to participate in C5aR-driven signaling but is not inhibited by FR and YM. Accordingly, we cannot definitively rule out a small contribution of Gα_15_ to the C5a-dependent responses, however, we think this is unlikely because C5a-induced PLCβ-dependent Ca^2+^ increases have been shown to be fully pertussis toxin (PTX) sensitive and mediated by Gβγ in this specific cell line (21).

Using immunolabeling coupled with three complementary fluorescence microscopy modalities (**Fig. S3A**), this work demonstrates that most of the PLCβ3 in macrophages (∼80%) is localized in the interior of the cell, away from the plasma membrane, which is consistent with previous results (18). Upon GPCR stimulation, PLCβ3 is recruited to the plasma membrane. Signaling from both Gα_q_ and Gα_i_-coupled receptors induced recruitment of PLCβ3 to the plasma membrane, indicating that both G proteins, Gβγ and Gα_q_, induce PLCβ3 membrane recruitment. In an earlier study using a reconstituted system Gα_q_ did not contain a lipid anchor and functioned solely to increase k_cat_ in PLCβ3-mediated PIP2 hydrolysis(6). By contrast, in Raw264.7 cells Gα_q_ also recruits PLCβ3 to the membrane. This difference likely reflects the presence of a lipid anchor on Gα_q_ in the cellular system and possibly differences in membrane lipid composition. The important point is, in the cellular system Gα_q_ is expected to both partition onto the membrane to recruit PLCβ3 and activate by increasing k_cat_.

Analysis of our TIRF images reveals an increase in PLCβ3 spot density as well as spot size upon stimulation with Gα_i_ or Gα_q_-coupled agonists. However, analysis of the super-resolution 2D STED images shows only an increase in PLCβ3 density without an effect of PLCβ3 spot diameter. These observations suggest that the larger spots observed in the TIRF images are a result of multiple distinct PLCβ3 enzymes within the diffraction limited (∼0.250 μm for green dyes) resolution of this approach, making the STED analysis more appropriate and robust to assess differences in membrane recruitment. In the STED analysis, stimulation of the Gα_i_-coupled receptor, C5aR, significantly increased the PLCβ3 membrane density at 10s, whereas stimulation of the Gα_q_-coupled receptor P2Y6 did not, suggesting that Gα_i_-coupled signaling may be more efficient at recruiting PLCβ3 to the membrane on the functionally relevant timescale in this context.

Our experiments demonstrate that Gα_q_ is not required for Gβγ-dependent activation of PLCβ, which is in agreement with other reports using cellular models (12, 13, 31). These observations are also supported by a significant body of work using *in vitro* experiments, which clearly demonstrate that Gβγ alone can robustly activate PLCβ enzymes (6-11). However, these results directly contradict conclusions from previous studies in other model systems showing that Gα_q_ is strictly required (14, 15, 17, 18). The biggest difference in these studies is the cell type and model systems used. We propose that the ability of Gβγ to independently activate PLCβ depends on the free Gβγ concentration resulting from receptor activation, which varies between cell types and activation stimuli (**Fig. 6**). Consistent with this proposal, a growing body of evidence suggests that the ability of Gα_i_-coupled receptors to regulate PLCβ is context dependent and is dictated by the local concentration of Gβγ generated by receptor stimulation (13, 20, 21) (**Fig. 6**). Similar observations were made regarding Gα_i_-coupled stimulation of GIRK channels (32, 33). The effective Gβγ concentration is dependent on several factors, including receptor density and organization, the kinetics of G protein release upon stimulation, and the Gβγ subtype and cellular distribution (20, 32, 33). Another factor likely contributing to this capability is the concentration of PLCβ on the plasma membrane at rest, which is proposed to be dictated by subtype and recruitment by additional proteins (1, 3). If sufficient PLCβ is not present on the plasma membrane, the concentration of Gβγ required to shift the equilibrium to membrane-associated PLCβ is higher (**Fig. 6**).

**Figure 6:**
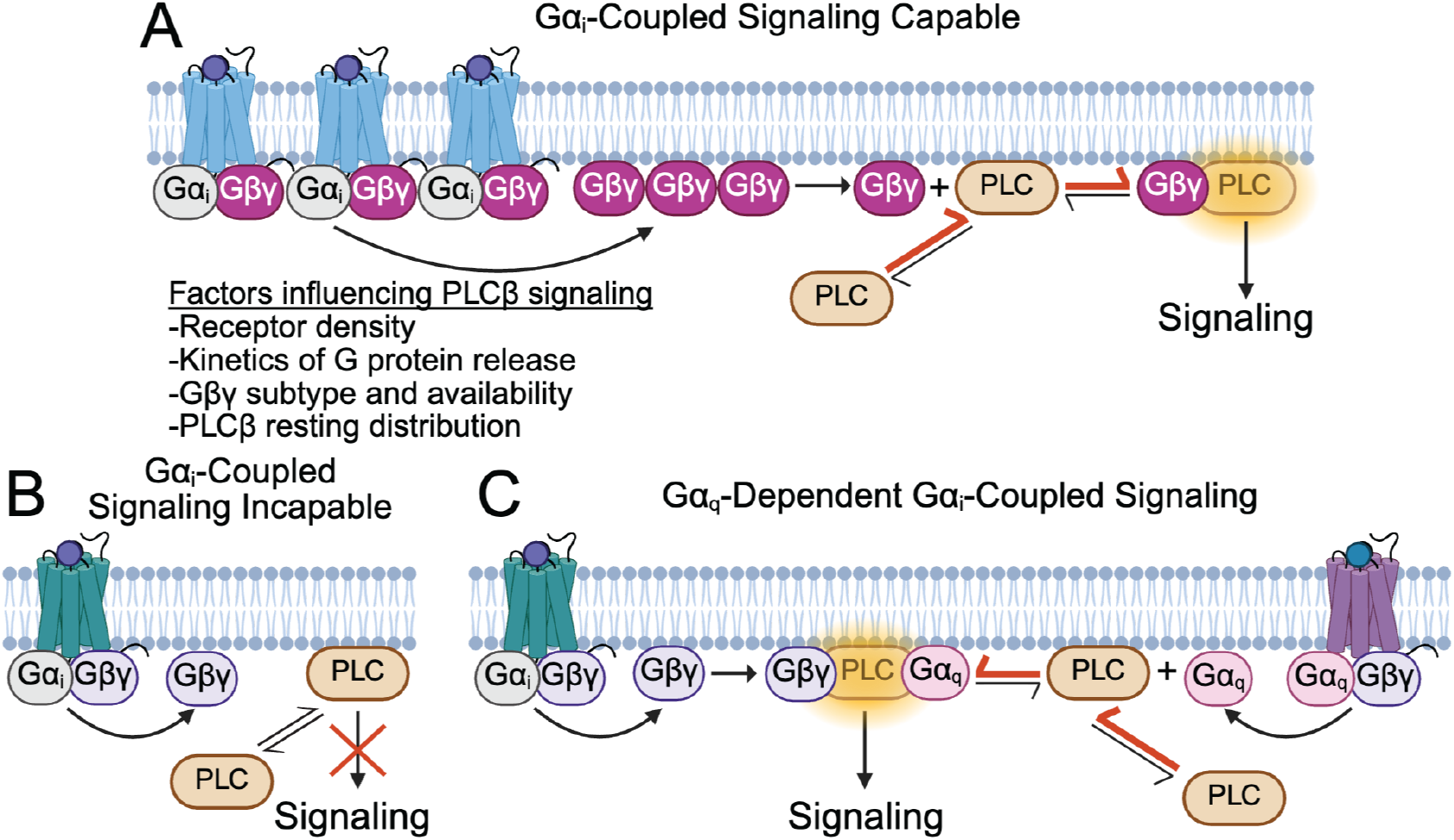
Gα_i_-induced PLCβ signaling is context dependent. **A:** In signaling capable contexts, the receptor density, kinetics of G protein release, Gβγ subtype and availability, and PLCβ resting distribution allow for sufficient Gβγ to be released upon receptor stimulation. This Gβγ recruits PLCβ to the membrane resulting in higher activity and thereby shifting its equilibrium from cytosolic to more membrane associated (red arrows). **B:** In signaling incapable contexts, the generated Gβγ is not sufficient to recruit PLCβ to the membrane so signaling does not occur. **C**: In Gα_q_-dependent contexts, released Gα_q_ recruits additional PLCβ to the membrane (red arrows) and also increases PLCβ catalytic rate (6, 7), rendering the previously insufficient Gβγ concentration able to promote signaling.

Such a concentration dependence is less relevant for Gα_q_-dependent stimulation because the affinity of PLCβ for Gα_q_ is much higher than for Gβγ, given that the PLCβ-Gα_q_ complex isstable in the absence of the membrane surface, which supports much higher local concentrations (3, 6). Additionally, our results, in agreement with other recent work (18), show that Gα_q_ signaling also recruits PLCβ to the plasma membrane, providing a mechanism for Gα_q_ to contribute to Gβγ-dependent activation, despite the observations that Gβγ-dependent activation can be independent of Gα_q_. In the case of stimulation with both Gα_q_- and Gα_i_-coupled agonists, Gα_q_ both allosterically activates and recruits PLCβ to the membrane. Thus, dual stimulation would reduce the concentration of Gβγ necessary to form PLCβ-Gβγ complexes where Gβγ can orient the PLCβ catalytic core for catalysis and further shift the PLCβ equilibrium from cytosolic to membrane-associated (**Fig. 6C**). A requirement for dual stimulation would make sense in physiological contexts where PLCβ stimulation would be detrimental to Gα_i_-coupled signaling, for example in GIRK signaling in cardiomyocytes (32, 34). Acetylcholine-induced activation of GIRK channels functions to induce hyperpolarization, which lengthens the interval between cardiac action potentials, thereby reducing heart rate (32, 34). This process is mediated by free Gβγ generated by Gα_i_-coupled acetylcholine receptors, but GIRK channels also require PIP2 to open (35, 36). Therefore, Gβγ-induced PLCβ activation would deplete PIP2 and counteract channel activation. This example highlights the necessity for intricate fine-tuning of PLCβ function, which requires additional layers, including rates of Gβγ release by receptors, the presence of competing Gβγ-binding proteins, and membrane localization of GPCRs and target proteins, to supplement the G protein-dependent regulation. While additional studies are needed to validate this proposal, it is an attractive hypothesis to explain inconsistencies in the literature regarding this system over the last two decades.

## Materials and Methods

### Cell Culture

Raw264.7 cells were obtained from ATCC (TIB-71) and were cultured according to ATCC recommendations. Briefly, they were maintained in Dulbecco’s Modified Eagle Medium (GIBCO) supplemented with 10% fetal bovine serum and 2 mM L-glutamine. For passaging, they were detached using a cell scraper. They were tested for mycoplasma during the course of the experiments.

### Compound stock concentrations

UDP (disodium salt, Tocris Bioscience 311150) was dissolved in H_2_O to make a 20 mM stock, which was diluted into DPBS +Ca^2+^/Mg^2+^ (Corning) for use in experiments. LPA (18:1 Lyso PA; Avanti Research 857130P-25mg) was dissolved in chloroform and the chloroform was evaporated using a continuous stream of Argon gas to produce a lipid film. The film was washed with pentane, followed by evaporation with Argon gas. The lipid film was rehydrated with H_2_O to make a 5 mM stock and sonicated to clarity with a bath sonicator. This stock was further diluted into DPBS +Ca^2+^/Mg^2+^ (Corning) for use in experiments. C5a was either purchased from abcam (Recombinant human C5a protein ab167724) or purified as described below. Commercial C5a was reconstituted with H_2_O at 36 μM and diluted into DPBS +Ca^2+^/Mg^2+^ (Corning) for use in experiments. U73122 (Tocris Bioscience 1268) and negative control U73343 (Tocris Bioscience 4133) was dissolved in chloroform, and the chloroform was evaporated with a continuous stream of Argon gas to produce a film, which was stored at -20°C until use. For experiments, the film was reconstituted in DMSO at 12 mM using a bath sonicator. FR900359 (FR) (caymen chemicals 33666) was dissolved in DMSO to make a stock of 10 mM. YM254890 (YM) (tocris YM254890) was dissolved in DMSO to make a 2 mM stock.

### C5a expression and purification

C5a was prepared as described (37). Constructs were synthesized with N-terminal Thioredoxin (Trx) and His tags followed by a 3C precession protease cleavage site and cloned into a Pet28 expression vector. SHuffle T7 Express Competent E. coli (NEB) were transformed according to the manufacturer’s protocol and incubated at 30°C overnight. Large cultures were grown to an OD_600_ of ∼1.0 and induced with 500 mM Isopropyl β-D-1-thiogalactopyranoside (IPTG) at 16°C overnight. Cells were harvested by centrifugation at 3,500 x g for 15 minutes and pellets were flash frozen and stored at -80°C until use. Purification was carried out at 4°C. Cells were resuspended in purification buffer (50 HEPES pH 7.5, 300 mM NaCl) supplemented with 30 mM imidazole, DNase and protease inhibitors (12.5 μg/mL leupeptin, 12.5 μg/mL pepstatin A, 625 μg/mL AEBSF, 1 mM Benzamadine, 100 μg/mL Trypisin inhibitor, 1x aprotinin, and 1 mM PMSF), lysed by sonication, and clarified by centrifugation at 39,000 x g for 45 minutes. Clarified lysate was bound in batch to 10 mL NiNTA resin (Qiagen) equilibrated with purification buffer supplemented with 30 mM imidazole for 1 hour. The resin was washed in batch with 100 mL of purification buffer supplemented with 30 mM imidazole then loaded onto a column and washed by gravity flow with 10 column volumes of high salt buffer (50 HEPES pH 7.5, 1 M NaCl, 30 mM imidazole) and eluted with purification buffer supplemented with 500 mM imidazole. Eluted protein was concentrated to ∼2 mL using a 15-mL Amicon concentrator with 10-kDa molecular weight cutoff, then diluted to ∼25 mL using purification buffer without imidazole and incubated with 3C precession protease (prepared in-house) overnight at 4°C remove the Txr and His tags. The cleaved protein was passed over a column of 10 mL NiNTA resin equilibrated with purification buffer supplemented with 35 mM imidazole three times to remove the free Trx-His and concentrated to 1 mL using a 15-mL Amicon concentrator with 3-kDa molecular weight cutoff. C5a was purified further via size exclusion chromatography using a Superdex 75 10/300 increase column equilibrated with purification buffer. Fractions containing C5a were pooled and concentrated to ∼0.8 mg/mL using a 4-mL Amicon concentrator with 3-kDa molecular weight cutoff and flash frozen and stored at -80°C until use.

### Ca^2+^ mobilization

For Ca^2+^mobilization experiments, Raw264.7 cells were seeded in black, clear bottom, poly-L-lysine coated 384 well plates (20 μL of cells at 0.7*10^6^ cells/mL) for at least 12-18 hours prior to the experiment. Cells were loaded with 20 μL of 2X Fluo-4 supplemented with 5 mM water soluble probenecid using the Fluo-4 Direct™ Calcium Assay Kit (Invitrogen) for 2 hours at room temperature protected from light. Fluorescence was monitored with excitation at 470 nm emission at 540 nm using a Hammamatsu μCell FDSS system with a sampling interval of 0.32 s. Agonists were automatically injected and mixed with continuous reading during the measurements. Final concentrations added were as follows: 10 μM UDP, 20 nM C5a, 5 μM LPA or 20 nM C5a+10 μM UDP.

For experiments with U73122 (20 μM), U73343 (20 μM), FR (10 μM), YM (500 nM), and DMSO (0.5%), compounds were diluted into the 2X Fluo-4 loading solution and incubated during the loading (2 hours at room temperature). DMSO was kept at or below 0.5% final concentration, which did not influence signaling responses (**Fig. S1**). PTX (Sigma, P2980-50UG) was added into the culture medium at 200 ng/mL for overnight incubation and added into the 2X Fluo-4 loading solution during the loading (2 hours at room temperature), for a final concentration of 300 ng/mL

Raw fluorescence traces were baseline subtracted and averaged over 8-12 wells for each experiment. For comparison, the peak heights from each individual trace were averaged to determine a final value for the peak height for each condition. The linear component of the peak increase was fit for each set of averaged traces with each technical repeat treated separately. Conditions were repeated at least 3 times with 2 technical replicates per biological replicate resulting in a total of n ≥ 6 averaged repeats.

### Immunolabeling

For immunolabeling, precision cover glass #1.5H (Thorlabs) coverslips were sterilized with ethanol, placed in the wells of a 24-well plate, and coated with poly-L-Lysine. Cells were seeded into wells with coverslips (0.5 mL of cells at 0.2*10^6^ cells/mL to ensure well-separated single cells for imaging) for 12-18 hours prior to the experiment.

All washes/incubations were 0.5 mL with gentle rocking unless otherwise noted. For immunolabeling, culture media was removed and cells were washed with DPBS containing Ca^2+^/Mg^2+^ (Corning, used throughout this procedure) for 5 minutes. Agonists for stimulation were diluted in DPBS at the following concentrations: 10 μM UDP, 20 nM C5a, or 20 nM C5a+10 μM UDP. Agonists were added and incubated for the desired stimulation time and fixing was initiated with 8% PFA for 5 minutes with gentle rocking. The solution was exchanged for DPBS+4% PFA and incubated for 10 minutes. Cells were washed twice with DPBS for 5 minutes and permeabilized with DPBS supplemented with 2% BSA (DPBS+BSA) and 0.5% Triton X-100 for 10 minutes. The Triton X-100 was washed out with DPBS+BSA and incubated for 20 minutes. Cells were washed an additional time with DPBS+BSA and incubated with primary antibodies (see **Table 1**) for 1 hour. The cells were washed 3 times with DPBS+BSA for 5 minutes each and incubated with secondary antibodies (see antibodies table) for 1 hour protected from light. The cells were washed 3 times with DPBS+BSA for 5 minutes each and re-fixed with DPBS+BSA supplemented with 4% PFA for 10 minutes. The PFA was washed out twice with DPBS+BSA and coverslips were mounted on glass slides with ProLong diamond antifade mountant (Invitrogen) and sealed immediately with nail polish to prevent curing, 3D flattening and to retain a lower refractive index required for TIRF microscopy. Slides were stored to 4°C and imaged within a week of preparation. Stimulation time courses were completed with two biological repeats with two technical repeats for each biological repeat.

For experiments in the presence of PTX and FR treatments, cd11b was used as a marker for the plasma membrane. Cells were plated overnight with PTX (300 ng/mL) or for 2 hours with FR (10 μM) as described for the functional experiments. C5a stimulation and staining for PLCβ was carried out as above. To stain for cd11b, cells were re-fixed with DPBS+BSA supplemented with 4% PFA for 10 minutes and the PFA was washed out twice with DPBS+BSA, followed by incubation with Alexa Fluor® 594 labeled primary antibody against cd11b for 1 hour protected from light. The cells were washed 3 times with DPBS+BSA for 5 minutes each and re-fixed with DPBS+BSA supplemented with 4% PFA for 10 minutes. The PFA was washed out twice with DPBS+BSA and coverslips were mounted as described above.

### TIRF imaging

TIRF imaging was performed using an inverted widefield fluorescence microscope stand (Nikon ECLIPSE Ti2) equipped with motorized XY stage, Perfect Focus System (PFS), and a iLas 2 (Gataca Systems) illumination device allowing Azimuthal TIRF imaging (ring-TIRF), a specialized TIRF microscopy technique designed to improve illumination uniformity of evanescent wavefront by reduction of interference artifacts. This system allows highly calibrated imaging depth control from widefield epifluorescence, to Highly Laminated Optical Sheet imaging (HILO, an oblique illumination technique), to TIRF imaging down to ∼70 nm shallow depth of field. The system is equipped with 488/561/640 nm diode lasers and appropriate excitation and emission filters for 3-color TIRF imaging. 1200X1200 pixel frames of images were acquired through a CFI Apochromat TIRF 100XC NA 1.49 Oil objective lens (Nikon) with motorized correction collar on a Orca Fusion sCMOS camera (Hamamatsu) with 6.5 µm pixel pitch, using NIS Elements v5.42.06 (Nikon) software suite. Images were saved as the system default “.nd2” file format that retains all image metadata.

### STED imaging

STED imaging was performed on an inverted IX81 microscope stand (Olympus, now Evident Scientific), equipped with motorized X,Y stage, Zero Drift Compensation hardware based autofocus (ZDC2), and a Facility Line STED system (Abberior) with a pulsed 775nm depletion laser and 405nm, as well as pulsed 488/561/640 nm excitation laser lines. Spatial Light Modulators (SLM) generated specific 2D/3D STED depletion beam patterns that were delivered to the stand via a single beam path with a deformable mirror adaptive optical accessory to correct for refractive index mismatch and spherical aberrations. Images were acquired using Imspector LightBox software v. 16.3.14287-w2129 (Abberior), and an UPLXAPO100X 1.45 NA oil immersion objective (Olympus, now Evident Scientific) and 3D STED was performed using a mix of 90% 3D and 10% 2D depletion patterns, while keeping the pinhole closed to ∼0.7 AU. Images were acquired on separate spectrally gated Avalanche Photodiode detectors (APDs) using 650-755 nm band pass for STAR RED and 588-698 nm band pass for Alexa Fluor® 594. Image pixel/step sizes were calculated to satisfy the Nyquist sampling criteria using an online calculator tool (https://svi.nl/Nyquist-Calculator), and all images were collected using the same settings for excitation and depletion laser powers, pinhole diameter, line accumulation and pixel dwell times. Acquired images were saved as the default “.obf” files that retain all image metadata.

### Image processing and analysis

Images were processed and analyzed using Huygens Professional v25,10 (SVI). Templates were generated for microscopy parameters based on image metadata, and for deconvolution using an appropriate point spread function (PSF) model for automatically computing a theoretical PSF from microscopic parameters and a model of the microscope. All images were batch processed for consistency using the templates in Huygens Workflow processor. The deconvolved images were further corrected for chromatic aberrations and analyzed using Huygens Object Analyzer advanced module. Appropriate thresholds and seeds for intensity masks were determined empirically at first, with automatic Otsu’s thresholding as a starting point, and the segmentation quality were fine-tuned further using Gaussian filtering with a low sigma radius and watershed fragmentation with sparse seeding. All comparable images were segmented with identical parameters and measurements were extracted for the reported parameters as “.csv” files for further analysis.

For TIRF images, full image frames were cropped to individual cells and subjected to express deconvolution using Huygens Professional before processing and analysis as described above. Within the object analyzer, cell boundaries were manually traced using the C5aR signal for cells not treated with PTX/FR and cd11b for cells treated with PTX/FR and the PLCβ spots/area and spot diameter were measured. At least three images for each technical repeat were collected, resulting in n=12 images/time point. At least two cells were cropped from each image, resulting in n>24 cells/time point in the final statistics shown in **Fig. 3 and S4**.

For 3D STED axial line scans, the plasma membrane was manually traced using the C5aR signal and the percentage of PLCβ puncta within 100nm of this tracing was measured. At least 4 images for each technical repeat were collected, resulting in n>16 for each condition in the final statistics in **Fig. 4**.

For 2D STED analysis, images were cropped in z to include only the plasma membrane proximal part of the Z-stack using the C5aR signal as a guide. Density of C5aR and PLCβ spots was determined within the cropped image volume, and the X-Y diameter of each spot was reported. To ensure the results were not dependent on the thresholding used, results for two different segmentation parameters are reported (**Fig. S5**). At least three images for each technical repeat were collected, resulting in n ≥ 6 images for each condition in the final statistics in **Fig. 5** and **S5-6**.

Two-sample Student’s t-test were completed with a null hypothesis that the difference between the means is 0 to assess the statistical significance of each time point from the nonstimulated condition. P values are indicated directly on the graphs.

## Acknowledgments

All microscopy work was performed in the Bio-Imaging Resource Center at Rockefeller University, RRID:SCR_017791. We thank Alison North and other members for their assistance with fluorescence imaging experiments and data analysis. We thank Chloe Larson and other members from the Fisher Drug Discovery Resource Center of the Rockefeller University for help and resources to complete the Ca^2+^ mobilization experiments. We thank members of the MacKinnon lab, Jue Chen, and members of her research group for helpful discussions. This work was supported by National Institute of General Medical Sciences (NIHF32GM142137 to MEF). RM is an investigator in the Howard Hughes Medical Institute. Some figures were created using BioRender.com.

## Data, Materials, and Software Availability

All study data are included in the article and/or supporting information.

