## Supplementary material for "PLCβs are recruited to the plasma membrane in macrophages by both Gβγ and Gα_q_": Figure S


**This PDF file includes:**

Figures S1 to S6

**Figures**

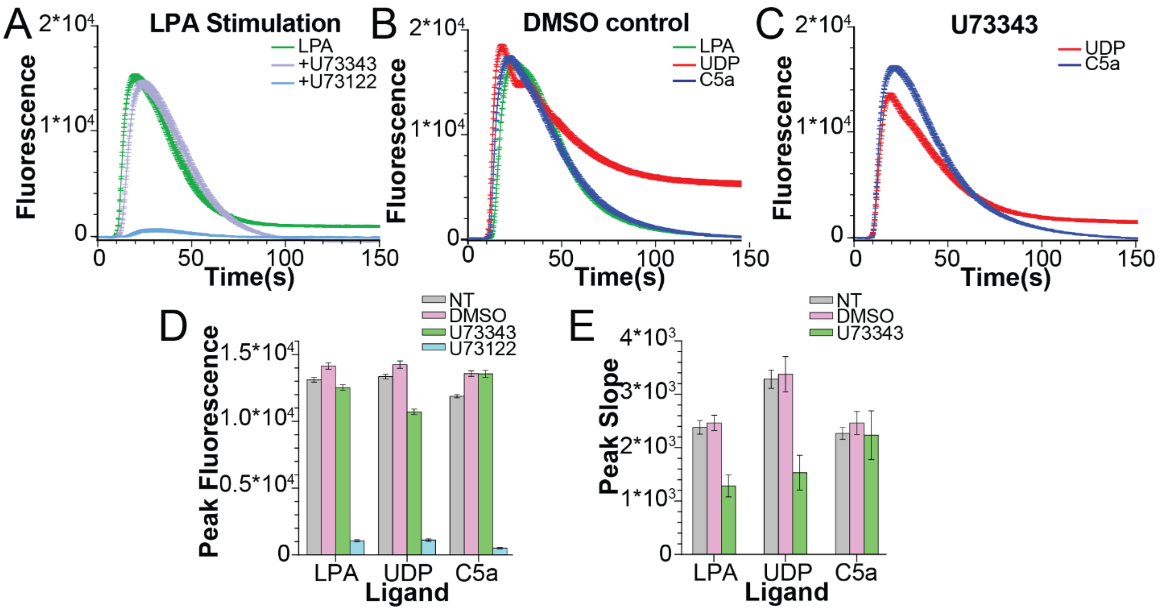


**Figure S1**: LPA-induced PLCβ signaling and U73343 validation. **A**: Representative $Ca^{2+}$ transients upon stimulation with LPA (green), LPA + U73343 (purple), and LPA + U73122 (light blue). **B-C**: Representative $Ca^{2+}$ transients upon stimulation with LPA (green), UDP (red), or C5a in the presence of 0.5% DMSO (B) or 20 μM U73343 (C). **D-E**: Quantification of $Ca^{2+}$ peak fluorescence (D) or slope of $Ca^{2+}$ peak (E) for LPA, UDP, and C5a in the presence of DMSO, U73343, or U73122. Error bars are SEM. For A-C, traces are averaged from 10 wells and error bars are SEM. UDP was added at 10 μM, LPA at 5 μM, and C5a at 20 nM final concentrations.

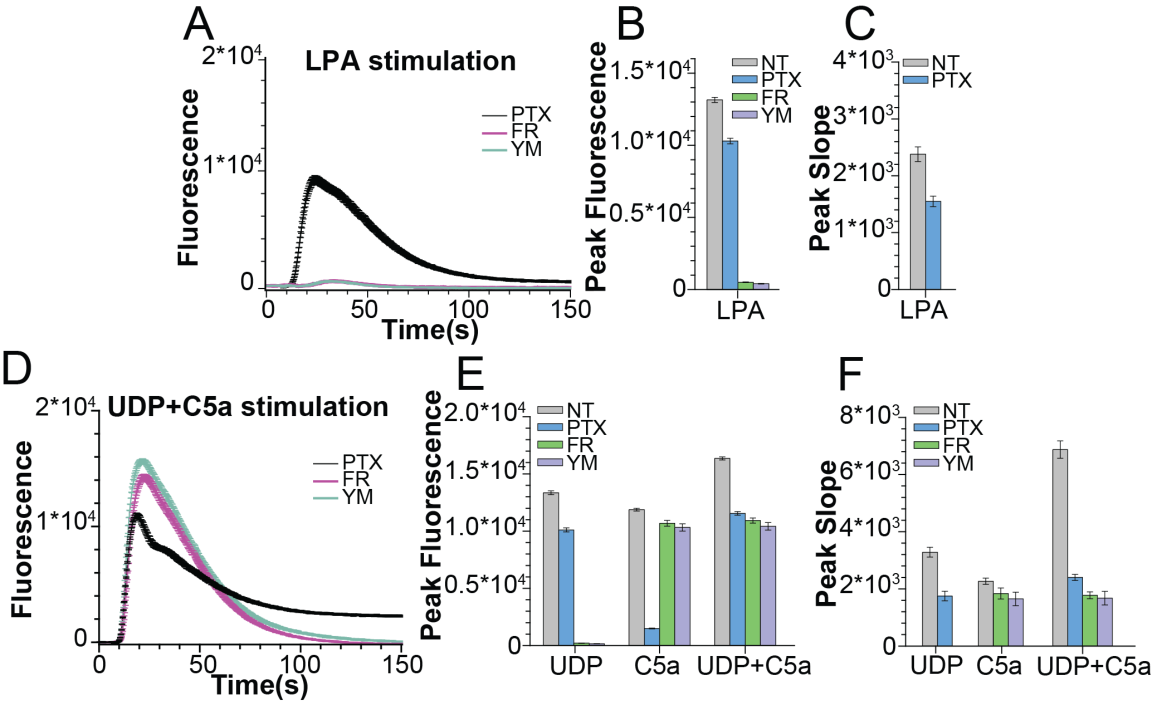


**Figure S2**: G$\alpha_q$ vs G$\alpha_i$ coupling of agonists. **A**: Representative $Ca^{2+}$ transients upon stimulation with LPA in the presence of 300 ng/mL PTX (black), 10 $\mu$M FR (pink), or 500 nM YM (light blue). **B-C**: Quantification of $Ca^{2+}$ peak fluorescence (B) or slope of $Ca^{2+}$ peak (C) for LPA in the presence of PTX, FR, and YM. **D**: Representative $Ca^{2+}$ transients upon stimulation with UDP+C5a in the presence of PTX (black), FR (pink), or YM (light blue). **E-F**: Quantification of $Ca^{2+}$ peak fluorescence (E) or slope of $Ca^{2+}$ peak (F) for UDP+C5a stimulation in the presence of 300 ng/mL PTX, 10 $\mu$M FR, and 500 nM YM. Error bars are SME. For A and D, traces are averaged from 10 wells and error bars are SEM. UDP was added at 10 $\mu$M, and C5a at 20 nM final concentrations.

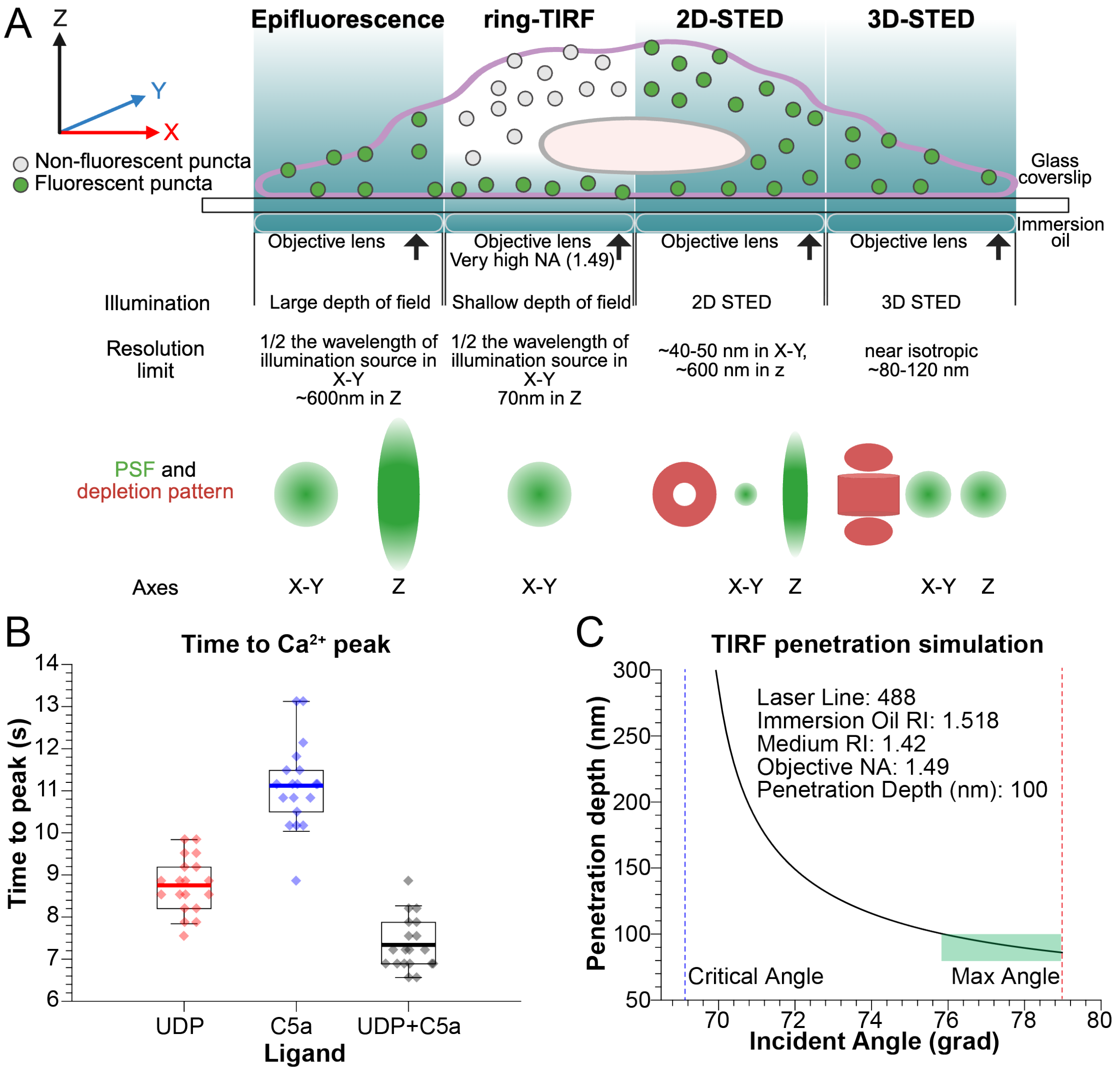


**Figure S3**: Microscopy Approaches. **A**: Summary of microscopy approaches. TIRF microscopy, which is diffraction limited, was used to analyze PLCβ3 spots at the plasma membrane at whole cell level. 2D STED, which provides very high 2D (X,Y) resolution, was used to obtain super-resolved information about the PLCβ3 puncta at the plasma membrane. 3D STED, which provides isotropic super-resolution, was used to study the fraction of PLCβ3 present at the plasma membrane. **B**: Quantification of time to $Ca^{2+}$ peak following stimulation with UDP (red), C5a (blue), or both (gray). Boxes are 25%-75%, error bars are 5%-95%, thick bar is the mean. Diamonds are individual experiments. **C**: Simulation using Calc TIRF depth plugin in ImageJ (https://imagej.net/ij/plugins/tirf/index.html) to model the penetration depth of evanescent light wave using our experimental parameters. Input parameters are shown in the table on the graph. The blue line shows the critical angle, demonstrating that TIRF was achieved in our experiments. The green box highlights the region of the curve with less than 100 nm depth, confirming that our TIRF images correspond mostly to the plasma membrane proximal cell regions.

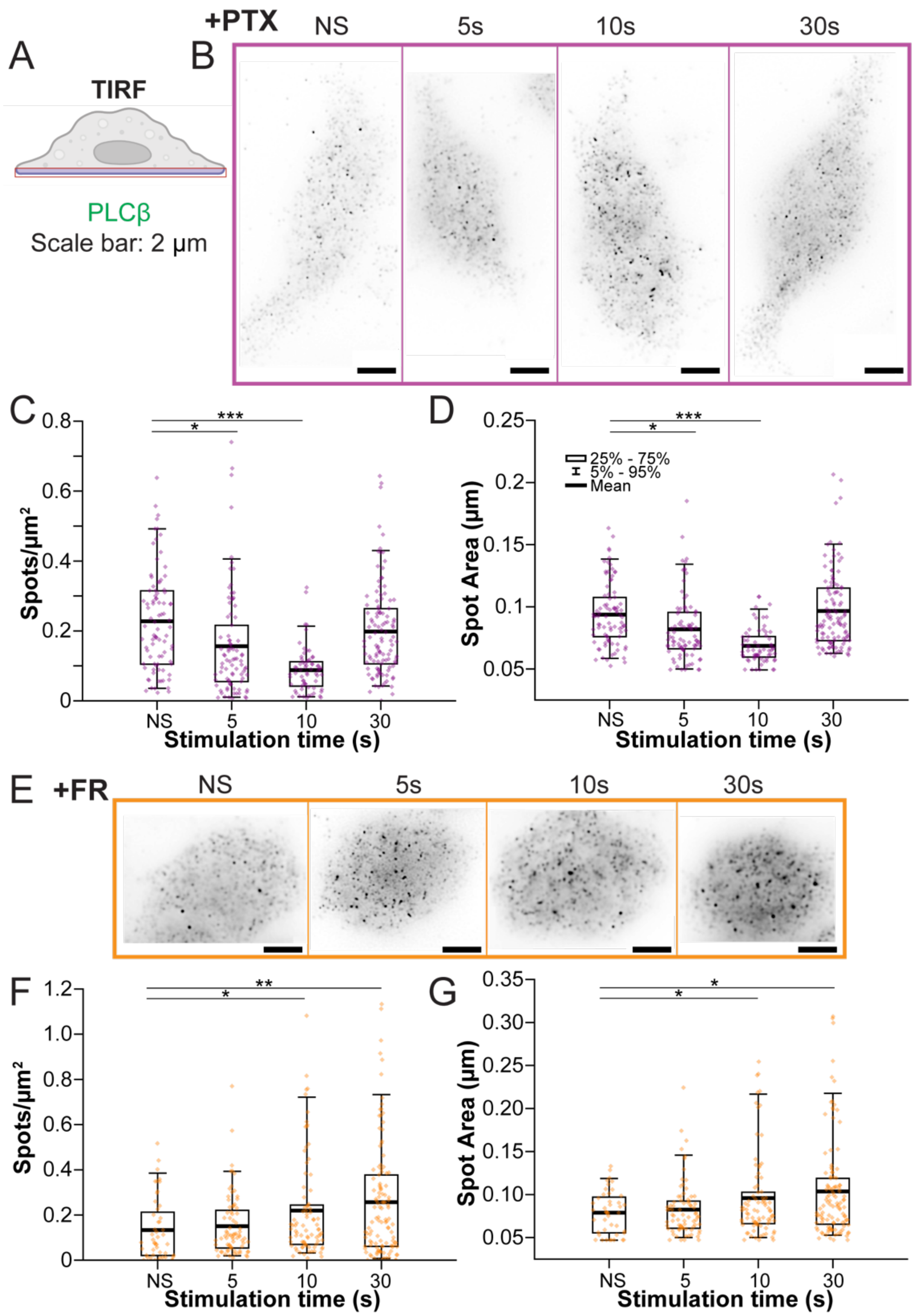


**Figure S4**: GPCR signaling recruits PLCβ3 to the plasma membrane. **A**: Cartoon of the TIRF imaging plane. **B,E**: Representative images of cells stained for PLCβ3 treated with PTX (B) or FR (E) prior to stimulation with 20 nM C5a. Images are shown with the AF488 channel LUT inverted to highlight cell boundaries using the diffuse cellular signal of AF488. Scale bar is 4 µm. Time points indicate the time of agonist exposure prior to cell fixation. **C,F**: Quantification of PLCβ3 spot density over time upon stimulation with C5a in the presence of PTX (C) or FR (F). **D,G**: Quantification of PLCβ3 spot size over time upon stimulation C5a with in the presence of PTX (D) or FR (G). NS is nonstimulated. Each point is a single cell. *: 0.05>p>0.005. **: 0.005>p>0.0005, ***: p<0.0005.

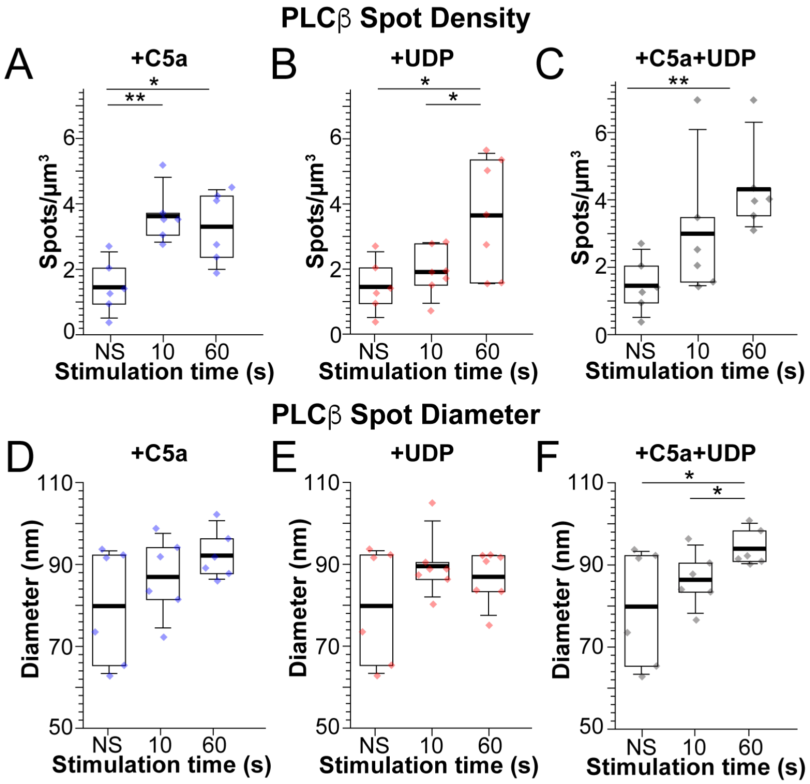


**Figure S5**: Analysis of 2D STED images with low threshold. **A-C**: PLCβ3 spot density on the plasma membrane over time upon stimulation with C5a (A), UDP (B), or UDP+C5a (C). **D-F**: Quantification of PLCβ X-Y diameter over time upon stimulation with C5a (D), UDP (E), or UDP+C5a (F). UDP was added at 10 μM and C5a at 20 nM final concentrations. Time points indicate the time of agonist exposure prior to cell fixation. NS is nonstimulated. Boxes are 25%-75%, error bars are 5%-95%, thick bar is the mean. Each point is from a single z stack. *: $0.05>p>0.005$. **: $0.005>p>0.0005$, ***: $p<0.0005$.

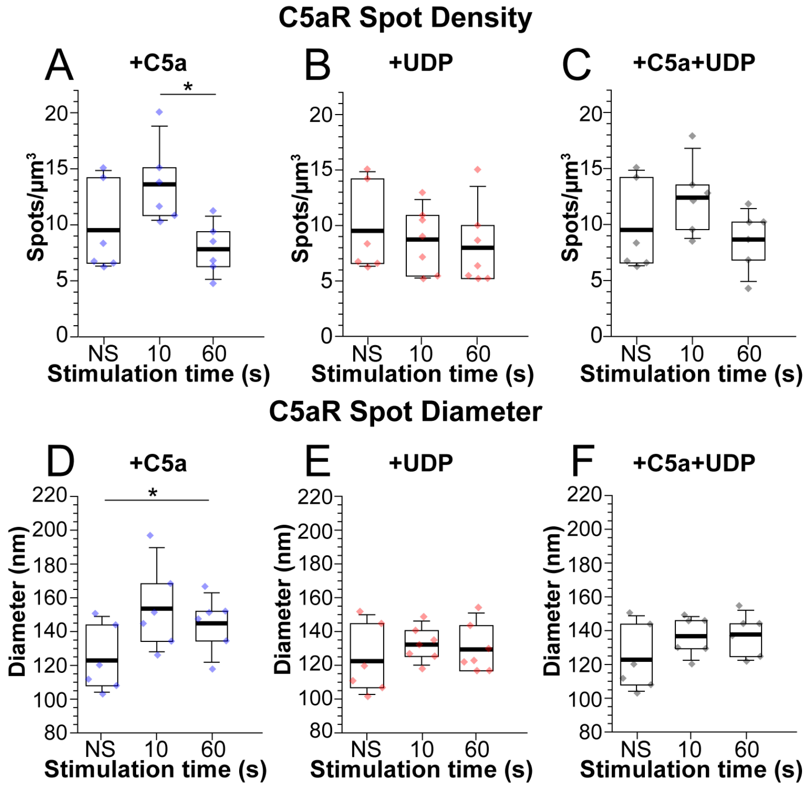


**Figure S6**: 2D STED analysis for C5aR. **A-C**: C5aR spot density on the plasma membrane over time upon stimulation with C5a (A), UDP (B), or UDP+C5a (C). **D-F**: Quantification of C5aR X-Y diameter over time upon stimulation with C5a (D), UDP (E), or UDP+C5a (F). UDP was added at 10 μM and C5a at 20 nM final concentrations. Time points indicate the time of agonist exposure prior to cell fixation. NS is nonstimulated. Boxes are 25%-75%, error bars are 5%-95%, thick bar is the mean. Each point is from a single z stack. *: $0.05>p>0.005$. **: $0.005>p>0.0005$, ***: $p<0.0005$.
